# On the impact of chemo-mechanically induced phenotypic transitions in gliomas

**DOI:** 10.1101/476150

**Authors:** Pietro Mascheroni, Juan Carlos Lopez Alfonso, Maria Kalli, Triantafyllos Stylianopoulos, Michael Meyer-Hermann, Haralampos Hatzikirou

**Affiliations:** Braunschweig Integrated Centre of Systems Biology and Helmholtz Center for Infectious Research, Braunschweig, Germany; Cancer Biophysics Laboratory, Department of Mechanical and Manufacturing Engineering, University of Cyprus, Nicosia, Cyprus; Centre for Individualised Infection Medicine, Hannover, Germany; Institute for Biochemistry, Biotechnology and Bioinformatics, Technische Universität Braunschweig, Braunschweig, Germany

## Abstract

Tumor microenvironment is a critical player in glioma progression and novel therapies for its targeting have been recently proposed. In particular, stress-alleviation strategies act on the tumor by reducing its stiffness, decreasing solid stresses and improving blood perfusion. However, these microenvironmental changes trigger chemo-mechanically induced cellular phenotypic transitions whose impact on therapy outcomes is not completely understood. In this work, we perform experiments to analyze the effects of mechanical compression on migration and proliferation of two glioma cell lines. From these experiments, we derive a mathematical model of glioma progression focusing on cellular phenotypic plasticity. The model reveals a trade-off between tumor infiltration and cellular content as a consequence of stress-alleviation approaches. We discuss how these findings can improve the current understanding of glioma/microenvironment interactions, and suggest strategies to improve therapeutic outcomes.

## Introduction

Gliomas originate from glial cells or their precursors in the brain and constitute the most common type of malignant brain tumor in adults^1,2^. The World Health Organization classifies gliomas into different categories, ranging from low-grade to high-grade malignancies^3^. The classification is primarily based on the tumor proliferative capacity and invasive tendency, with Glioblastoma Multiforme (GBM) being the most malignant, and unfortunately the most common, glioma in adults^2^. Notably, high grade gliomas are characterized by extensive, diffuse infiltration of cancer cells into the host brain tissue. Such aggressive behaviors are generally associated with poor patient prognosis and, for the case of GBM, almost 95% fatality rate within five years from diagnosis^4^. In fact, the infiltrative nature of gliomas is considered one of the main reasons for poor treatment outcomes. Despite recent progresses in neuro-oncology and advances in medical imaging, complete tumor eradication is often unsuccessful, leading to tumor recurrence in most of the cases. A better understanding of the mechanisms underlying glioma progression is urgently needed to develop effective treatments, and reduce glioma associated mortality.

In the last few years, it has become increasingly clear that tumor microenvironment plays a critical role in tumor progression and resistance to therapies^5–7^. Two important components characterize the tumor microenvironment, namely chemical and mechanical factors. The chemical microenvironment of the tumor comprises soluble factors released and uptaken by tumor cells. A major role among these chemicals is played by oxygen, and particularly by its deficiency. Indeed, hypoxia correlates with tumor invasiveness and malignancy^8^. Tumor cells respond to hypoxia by secreting agents that stimulate blood vessel formation, via *tumor angiogenesis*^9^, and activating cell migration^8^.

Regarding the physical environment, the growth of a tumor in the confined space of a host tissue is known to generate mechanical forces^10,11^. These forces hinder cell proliferation, induce apoptosis and enhance the cellular invasive potential of tumor cells^12–14^, as shown in the experiments discussed in the Results section.

In addition, resulting mechanical stresses can induce occlusion of blood vessels, reducing tissue perfusion and contributing to tumor hypoxia^10^. In response to chemical and physical stimuli, tumor cells exhibit phenotypic changes. A prominent phenotypic plasticity mechanism is related to the mutually exclusive switching between the proliferating and migratory phenotypes. This phenomenon, also known as the Go-or-Grow mechanism, has been suggested to impact the invasive potential of gliomas^15–17^. H4 cells shown in the Results section are an example of a glioma cell line that follows this mechanism, modulating proliferation and migration in accordance to mechanical compression. In fact, the interplay between tumor behavior and microenvironmental stimuli is being actively investigated, and it is becoming the target of novel therapeutic strategies^18^. For instance, vascular normalization and stress-alleviation approaches have been investigated to treat solid tumors^19^. In particular, vascular normalization therapies are designed to restore the balance between pro- and anti-angiogenic factors, inducing a more functional tumor vasculature with the aim to improve drug delivery. On the other hand, stress-alleviation therapies propose to reduce mechanical stresses within the tumor to improve tissue perfusion and the action of chemotherapeutic agents. This is achieved by targeting specific components of the tumor extracellular matrix, such as collagen and hyaluronan. Indeed, depletion of these constituents by repurposing of approved anti-fibrotic drugs reduces tumor stiffness and mechanical stresses, improves tissue hydraulic conductivity and decompresses tumor blood and lymphatic vessels^20^. However, due to the complexity of tumor-microenvironment interplay, success rates of those normalization therapies are largely variable and still under intense investigation (see for instance ref.^21,22^ for the case of vascular normalization therapy).

Mathematical modeling is emerging as a complimentary methodology to aid biologists and clinicians in the understanding of tumor-host interactions^23,24^. Mathematical models are powerful tools to decipher the mechanisms underlying glioma infiltration and progression, as well as to suggest potential strategies for therapeutic interventions^25^. In addition, they help bridging the knowledge gap between microenvironmental changes and cellular plasticity, by testing hypotheses that would be difficult to experimentally prove. Different modeling approaches have been developed so far, with models based on discrete, continuous or hybrid descriptions of tumor-microenvironmental interactions. Regarding the continuous approach, one of the most adopted is the macroscopic diffusion model, proposed by Swanson and coworkers^26^. This model is based on a reaction-diffusion equation describing tumor cell density, taking into account cellular motility governed by heterogeneous diffusion coefficients or diffusion tensors computed from medical imaging^27^. However, this modeling technique does not usually take into account the generation of mechanical forces within the tumor tissue. This has been incorporated by extending the diffusion models to include linear momentum balance, adding the presence of an external body force describing the displacement of brain tissue by tumor cells^28,29^. Another modeling approach makes use of the theory of mixtures to account for mechanical effects in a sound mathematical framework^30–32^. Within this description, a tumor is modeled as a saturated medium comprising solid (e.g. tumor cells) and liquid (e.g. interstitial fluid) phases, subjected to mass, momentum and energy balances.

In this work, we propose a mathematical model describing the induction of phenotypic transitions in cancer cells by both chemical and mechanical cues. The model is based on the previous work in ref.^33^, in which the authors introduce a model for glioma invasion accounting for nutrient-driven phenotypic switching. The aim of this study is two-fold: (i) to complement the previous model in ref.^33^ with mechanical-driven phenotypic transitions, and (ii) to evaluate the effects of a stress-alleviation treatment on glioma progression and invasion. Models describing the application of stress-alleviation treatments already exist in the literature^34^; however, to the authors knowledge, the influence of cellular phenotypic transitions - resulting from these microenvironmental treatments - on tumor progression have not been investigated yet.

In this study, we assume that phenotypic changes are dictated by nutrient availability as well as by mechanical stresses, with tumor cells becoming more prone to migration and reduced proliferation at high compression states and low nutrient levels (and viceversa). We report on how cells with different sensitivities to mechanical stresses react to such microenvironmental stimuli, and investigate tumor response via two main observables, measuring the tumor infiltrative zone and total burden. Our analysis focuses first on mechanical driven phenotypic transitions, evaluating the effects of different levels of cellular mechanosensitivity on tumor behavior. Then, we study the exclusive effect of nutrient induced phenotypic transitions, and evaluate the influence of tissue stiffness on tumor growth and invasion. Finally, we analyze changes in tumor behavior to both chemical (nutrient) and mechanical (compression) stimuli at different levels of mechanosensitivity and variations in the ratio of nutrient-to-mechanical response. We conclude by discussing insights drawn from model simulations and propose new research directions for future studies.

## Results

### Effects of mechanical compression on glioma cells

We performed *in vitro* experiments to test the effects of mechanical compression on glioma cells from two cell lines. Results are reported in Figure 1, whereas the experimental procedure is described in the Materials and Methods section. We performed wound closure and proliferation assays to quantify glioma cell migration and proliferation in two different cell lines, i.e. H4 and A172 cells. H4 cells respond to compressive stresses by increasing migration and decreasing proliferation, in accordance with the Go-or-Grow mechanism. Compared to them, A172 cells are less mechanosensitive, not modifying their proliferation rate and even decreasing motility. As the latter cells derive from aggressive tumors (i.e. glioblastoma), low mechanosensitivity could arise as an adaptation to harsh environmental conditions.

**Figure 1.**
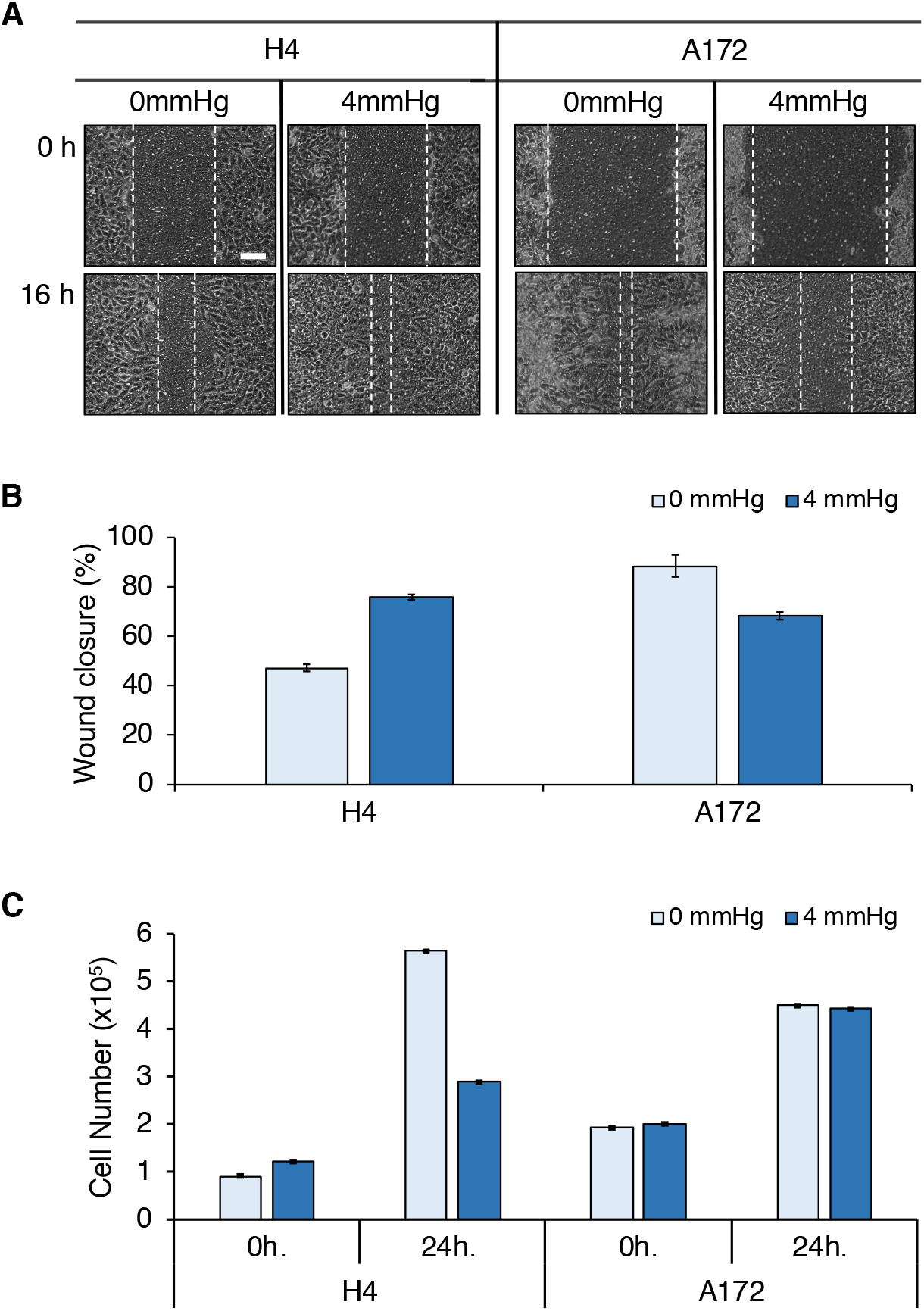
Different cell mechanosensitivities determine the response to physical cues. **A** Wound closure assay performed on two glioma cell lines, characterized by a more normal-like (H4) and metastatic (A172) behavior. Cancer cells are seeded on the inner chamber of a transwell insert, on the top of which an agarose cushion is positioned. A piston with adjustable weight applies a solid stress (4mmHg) on the cells and the effects of compression are visualized after 16h (scalebar: 0.1mm). **B** Quantification of wound closure at 16h. For H4 cells, compression increases migratory behavior, leading to a higher wound closure percentage. On the other hand, A172 cells display higher migratory properties in the control case (0mmHg), but respond to compression by reducing migration on the substrate. **C** Proliferation assay performed on H4 and A172 cells before application of mechanical compression (0, 4mmHg) and post compression (24h). Compression reduces proliferation in H4 cells, but it has negligible influence on A172 cell replication. In **B, C** error bars represent standard errors.

### Compression-driven phenotypic transitions

The mathematical model describes the growth of a vascularized tumor considering the interplay between two cell phenotypes, namely proliferative and migratory. The normalized system variables are the density of tumor cells *ρ*, the concentration of a nutrient, i.e. oxygen *n*, and the tumor vascular density *v*. Figure 2 shows a schematic representation of the system interactions and model assumptions, which are summarized below:

**A1** Glioma cells express a proliferative or migratory phenotype depending on the microenvironmental cues. In particular, high (low) nutrient availability and low (high) mechanical compression induce a proliferative (migratory) phenotype^8,12–17^.
**A2** Glioma cells consume nutrient provided by the vasculature^35^.
**A3** Increased mechanical pressure in regions of high glioma density induce blood vessel collapse and decrease nutrient availability^36^.
**A4** Oxygen is essential for glioma growth and progression^8,9^.
**A5** Blood vessels release nutrients^9,37^.
A6 The sprouting of new blood vessels stops when physiological levels of nutrients are restored^9,33^.

**Figure 2.**
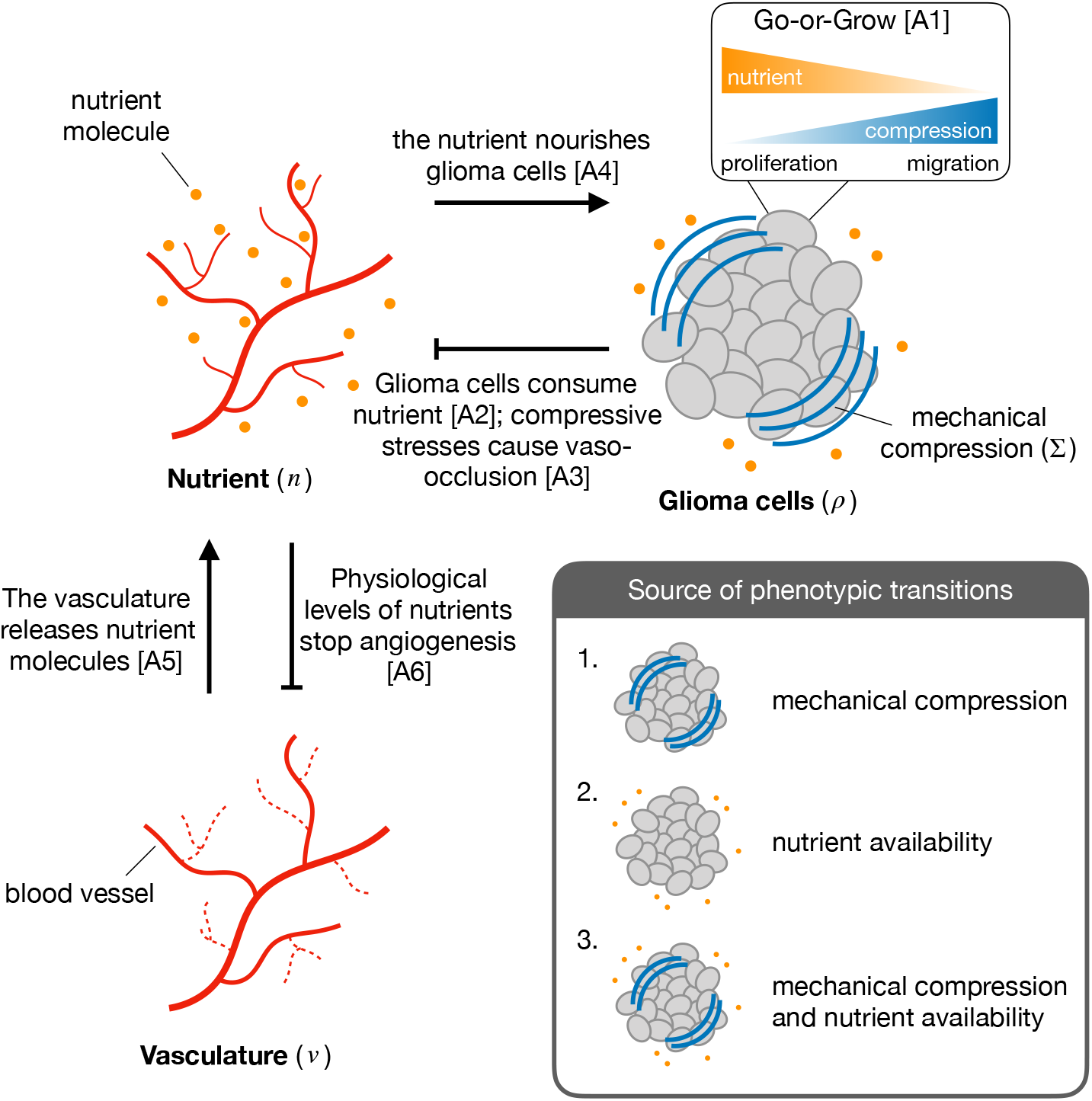
Scheme of the interactions between the different components of the system, i.e. glioma cells, nutrient availability and vasculature. The inset displays the different sources of phenotypic transitions in glioma cells considered in the model.

We characterize tumors by two quantities, namely the infiltration width (IW) and the tumor mass (TM), see the schematic in Figure 3A,B. Tumor IW is defined by the difference between the radial coordinates in which glioma cell density is 80% and 2% of the maximum cellular density, at the last time-step of simulations. In turn, the tumor mass is calculated by the integral

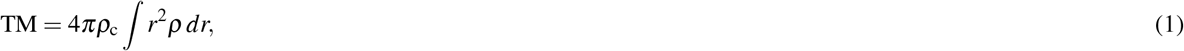

in which *ρ* is the tumor cell density at the last time-step of simulations, and *ρ_c_* is the carrying density of glioma cells. In the following sections, we investigate the dependence of IW and TM on different values of model parameters, as well as on different mechanisms driving cell proliferation and migration. Model results are presented as simulation maps showing the variations of the observable quantities over the (*D, r*) space, i.e. the parameter space of cellular intrinsic diffusion (*D*) and proliferation (*r*). Such space has been reported helpful for categorizing different glioma behaviors^38^. Indeed, by defining the non-dimensional number 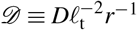, in which *ℓ*_t_ is the characteristic tumor length (≈ 1cm), it is possible to partition the pathophysiological (*D, r*) space in two distinct regions, see Figure 3C. In particular, for 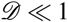, tumors are characterized by a dominant proliferative behavior, whereas for regions in which 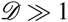 tumors display a more invasive behavior. Notably, the two major classes, i.e. low and high-grade gliomas, can be represented on the plane, occupying the bottom-left and top-right corners, respectively (see Figure 3C).

**Figure 3.**
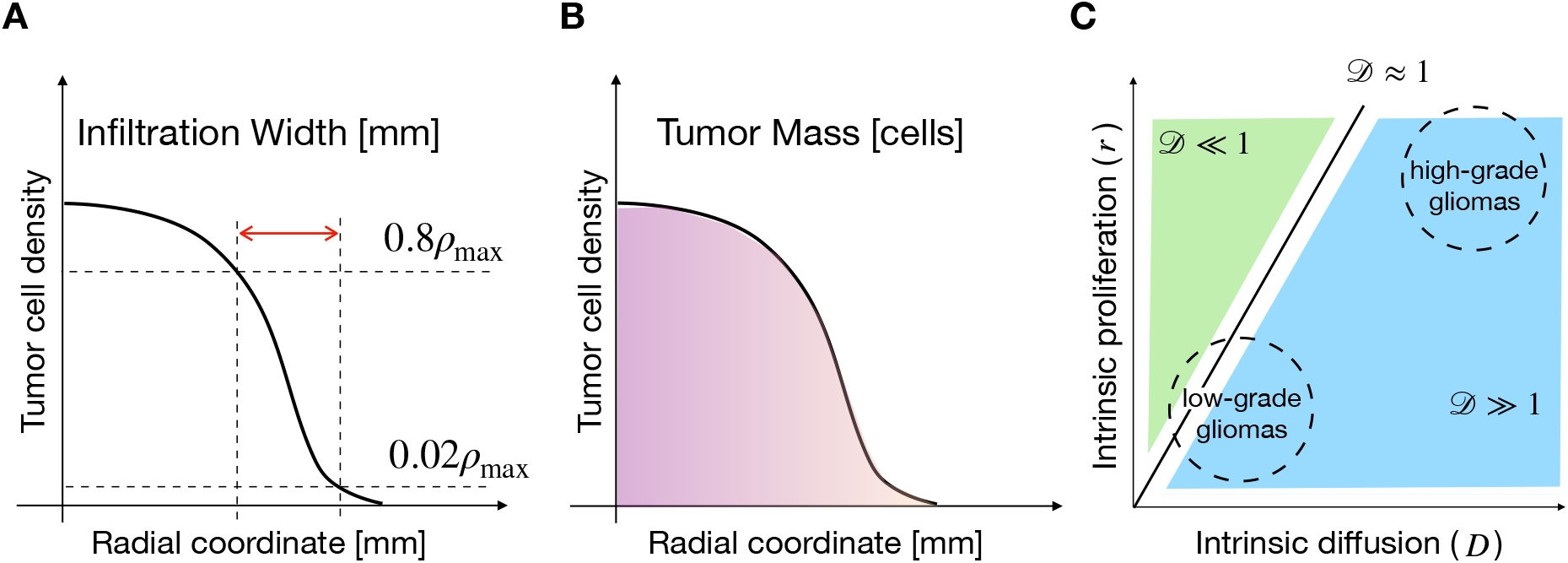
Graphical representation of model observables and glioma characteristics. Schemes representing tumor IW (**A**) and TM (**B**), describing tumor invasiveness and burden in the host, respectively. **C** Characteristics of gliomas across the cellular intrinsic diffusion and proliferation space. The relative importance of cell migration to proliferation is described by the non-dimensional number 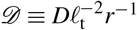, which splits the plane in regions where proliferation (green) or migration (blue) dominate. Low and high-grade gliomas occupy the bottom-left and top-right corners of the plane, respectively^38^.

We start our analysis by considering compression-driven phenotypic transitions. In this way, the evolution of the tumor is decoupled from the nutrient and vasculature equations - see the Materials and Methods section for a thorough derivation of the mathematical model. The tumor dynamics depend on *ασ*^−1^, a nondimensional number that represents the tissue stiffness effectively perceived by the tumor cells. For a fixed value of the tissue stiffness *α*, high values of *σ* (i.e. low mechanosensitivity) translate into poor cellular mechanical response. On the other hand, for low values of *σ*, tumor cells respond more intensely to mechanical compression. In the following, we will refer to *ασ*^−1^ as the *effective stiffness* of the tissue, and quantify its effect on tumor growth.

Figure 4A shows simulation maps of tumor IW and TM for different values of the effective stiffness. For increasing values of *ασ*^−1^, IW increases especially for highly-diffusive tumors. On the other hand, TM displays a non-monotonic trend for increasing effective stiffnesses. Indeed, we observe a maximum for this quantity at intermediate values of *ασ*^−1^. This finding is highlighted in Figure 4B,C where the maximum values of IW and TM are displayed for different *α/σ* ratios. Figure 4B shows that for low effective stiffnesses IW is short. However, for intermediate values of *ασ*^−1^, IW increases linearly with increasing effective stiffness. Moreover, for low values of *ασ*^−1^, tumor cells are not responsive to mechanical stresses and the tumor does not acquire an infiltrative behavior. On the other hand, for increasing values of *ασ*^−1^, the transition from proliferative to migratory phenotype is enhanced, leading to tumors with more diffusive fronts. The situation changes with respect to TM, where we observe a maximum for TM at intermediate values of *ασ*^−1^, see Figure 4C. This occurs because at low effective stiffnesses tumors are compact and their cellular density tends to the carrying capacity of the tissue, thus limiting the overall tumor growth. In the case of high *ασ*^−1^ values, tumor cells proliferate less and acquire instead an invasive phenotype, which reduces tumor burden.

**Figure 4.**
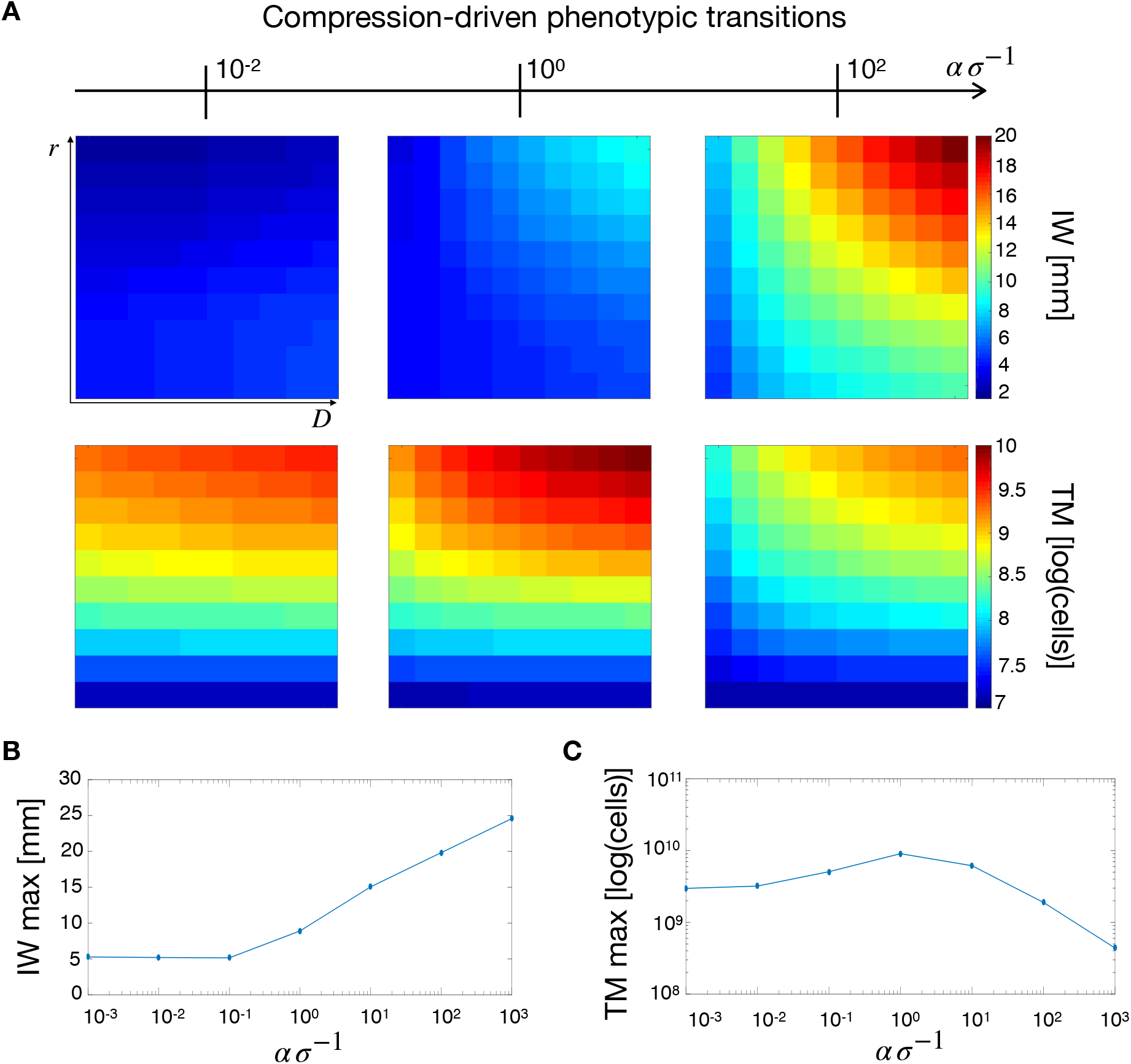
Tumor observables for mechanical-induced phenotypic switching. **A** Simulation maps of tumor IW and TM with respect to different values of effective stiffness *ασ*^−1^. From left to right *σ* = 10^5^, 10^3^, 10^1^Pa with *α* = 10^3^Pa. IW increases for increasing values of the *α/σ* ratio, whereas TM shows a maximum for intermediate values of effective stiffness. The maximum IW **B** and TM **C** are obtained for different values of effective stiffness, with *α* = 10^3^Pa, and *σ* varied. While IW shows an increasing trend with effective stiffnesses, TM displays an optimum for intermediate *ασ*^−1^ values.

### Nutrient-driven phenotypic transitions

Now we consider transitions between the two cell phenotypes driven only by nutrient availability in the tumor microenvironment. In this case, a critical role is played by the vascular response to mechanical stresses. If mechanical pressure overcomes a critical threshold blood vessels collapse and nutrients are not delivered to the tumor.

Simulation maps in Figure 5A show the influence of tissue stiffness *α* on IW and TM. As *α* increases due to tumor growth, intratumoral mechanical stresses are exacerbated. Thus, for higher tissue compliances (Figure 5A, left column), the overall IW map displays low values, with two peaks in the regions of low and high 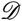. A similar behavior is observed for intermediate stiffness (Figure 5A, center column) where the peaks appear lightly shifted towards the region of higher 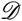. Finally, for high stiffness (Figure 5A, right column) IW displays a unique peak in the top-right corner, i.e. at high *D* and *r*. Indeed, when the stiffness is low and vascular occlusion is not engaged, the highest values of IW are observed in two extremes, for highly proliferating and highly diffusive tumors. Tumors with high values of *D* and *r* fall in the middle, with intermediate values of IW. The situation changes as soon as the vasculature collapses due to higher stiffness (and stress) and nutrient deficiency pushes the cells towards a more migratory phenotype. In this case, tumors with the highest intrinsic motilities and proliferation rates develop longer infiltrative fronts, leading to the peak in IW at the top-right corner of the simulation map in Figure 5. The situation changes for TM, (second row of Figure 5A), where even if the increase in stiffness does not produce alterations in the pattern of TM in the (*D, r*) space, it leads to an overall tumor burden reduction. This observation is in line with the deficiency of nutrients that follows from vascular occlusion at higher stiffnesses. As the tumor becomes more deprived of nutrients, cell proliferation is reduced and TM decreases.

**Figure 5.**
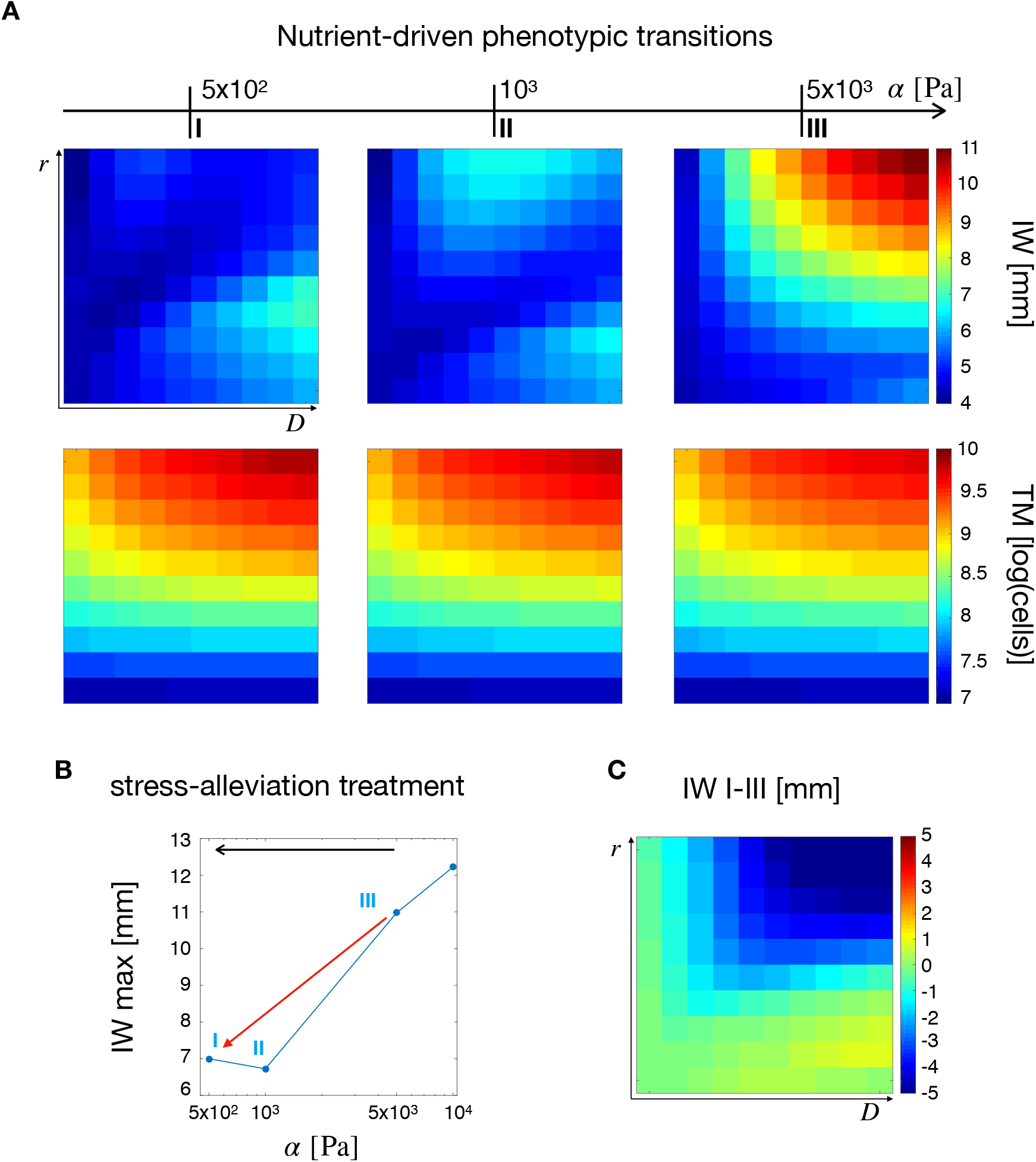
Tumor observables and the effects of a stress-alleviation therapeutic strategy in the case of nutrient-induced phenotypic switching. **A** From left to right, simulation maps of tumor IW and TM for varying tissue stiffness from 5 × 10^2^Pa to 5 × 10^3^Pa. A non-trivial effect of stiffness on IW and a monotonic decrease in TM for increasing values of *α* is observed. **B** Maximum value of IW for different tumor stiffness values. The red arrow points in the direction of a stress-alleviation treatment, where shorter IWs are obtained for decreasing values of *α*. **C** Simulation map for the difference in IW between the points I and III in B. Gliomas with the highest values of *D* and *r* display the highest reductions.

Figure 5B,C summarize the effects of tissue stiffness on tumor IW. In particular, Figure 5B shows a decrease in the maximum IW for decreasing *α*. This can be understood as a *stress-alleviation treatment*, in which the tumor is treated with matrix degrading agents to decrease tissue stiffness. In fact, a decrease in stiffness, and thus in mechanical stress levels, may lead to less malignant tumor phenotypes in terms of a reduction in IW^34^. This is shown in Figure 5C, in which the difference between IW at *α* = 5 × 10^2^Pa and *α* = 5 × 10^3^Pa is displayed over the (*D, r*) space. Simulations show a decrease of IW in the top-right corner of Figure 5C, which refers to the high-grade gliomas. Even though the nutrient and vasculature spatial profiles do not dramatically change in these regions (see Figure S1), the improvement in tissue perfusion is able to reduce the tumor invasive behavior.

### Chemo-mechanically induced phenotypic transitions

We now focus on the effects of both mechanical compression and nutrient deprivation on cellular phenotypic transitions. These are described by the ratio *t*_n_/*t*_s_, where *t*_n_ and *t*_s_ are the phenotypic transition rates related to nutrient availability and mechanical compression, respectively. For *t*_n_/*t*_s_ > 1 (*t*_n_/*t*_s_ < 1), cancer cells respond more (less) intensively to nutrient deprivation compared to mechanical stresses. We then simulate the effects of a stress-alleviation therapeutic strategy by decreasing the tissue stiffness from α = 5 × 10^3^Pa to α = 5 × 10^2^Pa. In addition, two ranges of effective stiffness are considered, namely *ασ*^−1^ = [10^−2^,10^−1^] and *ασ*^−1^ = [10^1^,10^2^], accounting for different degrees of cell mechanosensitivity.

#### Effects of phenotypic transitions on tumor IW

We first analyze the effects of cellular responses to microenvironmental stimuli on the tumor IW. Figure 6A shows the simulation maps for *t*_n_/*t*_s_ = 0.5 and *ασ*^−1^ = [10^−2^,10^−1^], i.e. in the case of mechanical response prevalent with respect to nutrient response and low mechanosensitivity. In the top row, simulations show higher IWs for increasing tissue stiffness, with the highest values at the top-right corner of the (*D, r*) space. The bottom row shows the effects of stress-alleviation treatments of different intensities, in which tissue stiffness is decreased from *α* = 10^3^Pa to *α* = 5 × 10^2^Pa (IW I-II, moderate intensity), and from *α* = 5 × 10^3^Pa to *α* = 5 × 10^2^Pa (IW I-III, high intensity). This results in only a slight variation of IW. Indeed, since in this case the cellular response to mechanical cues is stronger than for nutrients and the system is in the low mechanosensitivity regime, the effects of a stress reduction are limited on tumor IW.

**Figure 6.**
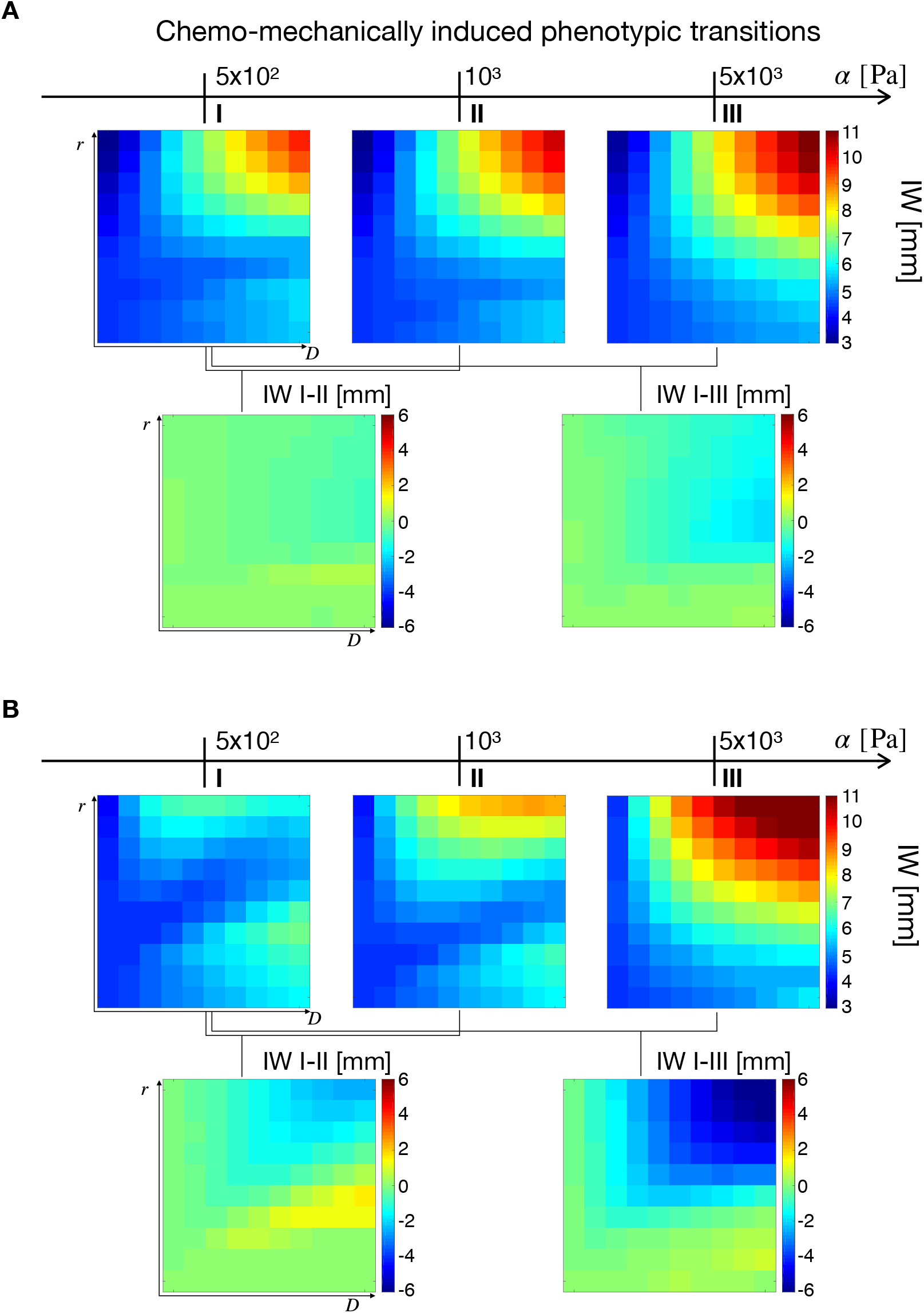
Simulation maps for the effects of chemo-mechanically induced transitions on tumor IW. In both cases (**A,B**), the top row shows three IW maps for different values of *α*, whereas the bottom row the IW variation occurring at different stiffness values. Simulations were obtained for *ασ*^−1^ = [10^−2^, 10^−1^] with *t*_n_/*t*_s_ = 0.5 (**A**) and *t*_n_/*t*_s_ = 10 (**B**).

Figure 6B deals with the case of nutrient response prevailing on the mechanical one, showing simulations with *t*_n_/*t*_s_ = 10 and *ασ*^−1^ = [10^−2^,10^−1^], i.e. low mechanosentivity. Results displayed in Figure 6B recapitulate the observations in the case of nutrient-driven transitions (Figure 5), with two peaks in the IW for low stiffness and a single maximum at high tissue rigidity. As the *t*_n_/*t*_s_ ratio takes large values it is reasonable to obtain similar results to the previous section (Figure 5), the mechanical response being quenched in the transition terms. Then, from the left map in the bottom row, two observations emerge. First, a slight decrease of IW in the top-right portion of the map is observed. This is a consequence of nutrient deprivation due to vascular occlusion, particularly occurring in high-grade tumors (high *D* and *r* values). Second, a slight increase of IW in the region of 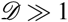 (as represented in Figure 3C), meaning that highly diffusive tumors benefit from a stress-alleviation strategy. The situation is different for the right simulation map in the bottom row of Figure 6B, referring to a more intense stress alleviation treatment (IW I-III). In this case, a decrease in IW for high-grade gliomas is obtained, similar to what can be observed in Figure 5B,C. As before, high-grade gliomas are more sensitive to changes in stiffness, and a reduction in mechanical compression results in a decrease of tumor IW.

The results for the IW difference at different *t*_n_/*t*_s_ ratios are summarized in Figure S2, in the Supplementary Material section. Simulation results show that by increasing the nutrient-to-mechanical response ratio the IW decreases in high-grade gliomas. This is particularly evident for the case of high intensity of stress-alleviation (displayed in the bottom row of Figure S2A). On the other hand, the top row shows a saddle in IW difference that emerges as *t*_n_/*t*_s_ increases. For high nutrient to stress response the model simulations qualitatively reproduce the results in ref.^33^. In that work, a biphasic variation of IW for high-grade gliomas for decreasing vaso-occlusion degrees was reported. In the case of low nutrient-to-mechanical response ratio, only slight variations of IW are observed for both intensities of the stress-alleviation strategy.

We perform the same analysis on nutrient-to-mechanical response in the case of high mechanosensitivity, i.e. *ασ*^−1^ = [10^1^, 10^2^]. For simplicity, we report the conclusive IW differences in the Supplementary Materials (Figures S2B, S3 and S4). For high *t*_n_/*t*_s_ ratios, the simulation maps resemble the results in Figure S2A, for low mechanosensitivity. Indeed, when the nutrient response overcomes the mechanical one, the tumor behavior is driven by nutrient availability and a decrease in IW occurs for high intrinsic cellular diffusion and proliferation. On the other hand, for low nutrient-to-mechanical response, a reduction of IW for all tumors represented in the (*D, r*) space is observed. Since tumor cells are more sensitive to mechanical stimuli, a reduction in the stress level translates in a less migratory cell population, and thus in a reduction of infiltrative patterns.

#### Effects of phenotypic transitions on TM

To conclude the study of chemo-mechanically induced phenotypic transitions, we analyze the effects of nutrient-to-mechanical response and mechanosensitivity level on TM. For the sake of brevity, we report the difference plots for TM in the various cases in the Supplementary Materials (see Figures S5, S6 and S7). Figure S5A shows the effects of phenotypic transitions on the variation of TM for different *t*_n_/*t*_s_ ratios at low cell mechanosensitivity. In this case, simulation maps show similar trends, with the intensity of the stress-alleviation treatment enhancing the variations in TM. For low *t*_n_/*t*_s_ ratios, a small decrease in tumor mass is observed at the top of the (*D, r*) space. This effect becomes less apparent when the nutrient response prevails over the mechanical one, see the simulation map for *t*_n_/*t*_s_ = 10. In this case an overall increase of TM is predicted for a significant part of the simulation map, with a particular increase for tumors with increased cellular motility and proliferation rate. Indeed, if *t*_n_/*t*_s_ < 1, a reduction in mechanical stress levels translates in a decrease of TM, as shown in Figure 4C for mechanical-driven transitions. Since cell mechanosensitivity is low, a reduction in the stress levels leads to dense tumors with a compact radius and a decreased TM. On the other hand, for high nutrient-to-mechanical response ratios, tumor evolution is significantly influenced by nutrient dynamics. When stress-alleviation strategies are pursued, vascular occlusion is reduced and nutrient supply is restored. This leads to tumors with higher densities and increased cellular proliferation.

To conclude the analysis for TM, the case of high mechanosensitivity, i.e. *ασ*^−1^ = [10^1^,10^2^], is investigated (see Figure S5B). In this case, the treatment induces a significant increase of TM for low *t*_n_/*t*_s_ ratios. Indeed, a predominant mechanical response in highly mechanoresponsive cells shifts the population to a more proliferative state in a stress-alleviation treatment. Cells turn from being for the most part migratory to actively proliferating, which results in a significant increase in TM. The increase is more evident for cells with high intrinsic proliferation rates, which contribute the most to the overall tumor burden. As the *t*_n_/*t*_s_ ratio increases, this behavior is gradually lost and the solutions for nutrient-driven phenotypic transitions are partially recovered (see Figure 5A). For *t*_n_/*t*_s_ = 10, an increase of TM is observed for high-grade gliomas, leading to tumors with reduced infiltrative patterns but increased cellular content.

#### High effective stiffness and low nutrient to stress response are the best fit for stress-alleviation treatments

Model results for the influence of stress-alleviation strategies on chemo-mechanically induced phenotypic transitions are summarized in Figure 7. In several cases the treatment produces opposing results: the IW decreases, but an increase in TM is observed at the same time. Indeed, if cells reduce their migratory phenotype due to the treatment, they become more proliferative, increasing TM. This may lead to tumors with high cellular densities but decreased invasive behaviors, possibly resulting in better outcomes for surgery. Another interesting point is visible in the bottom-left quadrant of the scheme, where the results for low mechanosensitivity and low nutrient-to-mechanical response are summarized. In this case the stress-alleviation strategy has limited impact on the tumor, being poorly effective in reducing its infiltrative behavior. For tumors in this area, adjuvant treatments potentially increasing cell mechanosensitivity may be sought, restoring the efficacy of the stress-alleviation strategy. To conclude, we estimate the impact of stress-alleviation strategies on the different cases by introducing a scoring system. We assign 0 to the cases in which treatment recommendation is low (L), i.e. the treatment does not produce significant changes or worsens tumor behavior. We assign 1 to the cases in which treatment recommendation is medium (M), e.g. reduction of IW together with increase of TM; and 2 for the cases of potentially successful outcomes (the treatment is highly recommended - H), i.e. reduction of IW and no change or decrease in TM. We report the labels of the different cases in Figure 7, and compute the resulting score for the fitness of the stress-alleviation treatment. The best fit to the treatment is obtained in the top-left quadrant, i.e. for low nutrient to stress response and high effective stiffnesses. On the other hand, low mechanosensitivity leads to poor scores, discouraging the use of this treatment for those situations.

**Figure 7.**
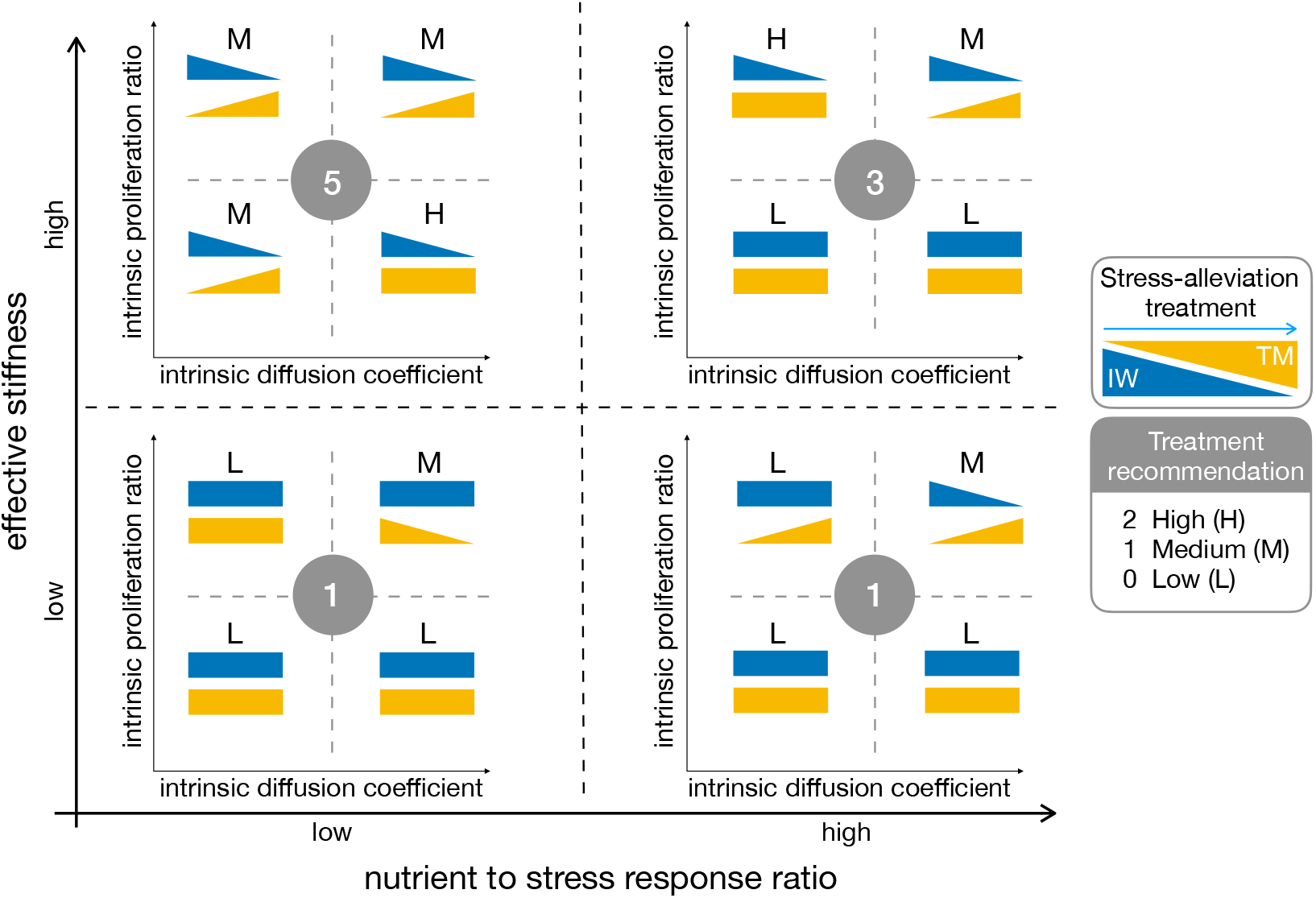
Summary of model results for chemo-mechanically induced phenotypic transitions. The triangles denote an increase (or decrease) of IW (blue) or TM (yellow) when a stress-alleviation treatment is performed. Rectangles denote a scarce effect of the treatment. The graph spans different levels of cell mechanosensitivity (vertical axis) and nutrient-to-mechanical response (horizontal axis). The labels and the circled numbers are related to the treatment recommendation score (L: low; M: medium; H: high), with higher numbers identifying better treatment outcomes.

Experiments such as those proposed in the Results section could be used to rank glioma cells for their intrinsic proliferation, migration and mechanosensitivity. Indeed, scratch assays provide information on cell motility, whereas cell duplication rate is obtained from proliferation assays. In addition, changes in cell proliferation and migration following mechanical compression are a proxy for the sensitivity of the cell line to mechanical stimuli. For instance, the H4 cell line is highly mechanosensitive and therefore a tumor including H4-like cells would highly benefit from the stress-alleviation treatment. On the contrary, an A172-like tumor would be non-plastic and stress-alleviation would not confer great advantages to the patient.

## Discussion

We presented a mathematical model describing tumor growth and invasion, and we specialized it to the case of gliomas. The model is based on a system of partial differential equations, coupling tumor, nutrient and vasculature dynamics. We focus on the Go-or-Grow mechanism, defining two cellular populations with mutually exclusive characteristics, i.e. migratory and proliferative cells. We analyzed the impact of microenvironmental stimuli, in terms of mechanical compression and nutrient availability, on glioma progression. In addition, we investigated the effects of cellular mechanosensitivity and nutrient-to-mechanical cellular response on two main tumor characteristics, namely the infiltration width (IW) of the tumor and its mass (TM).

Model simulations show that the combination of responses from external stimuli is able to generate a wide pattern of tumor behaviors. When the tumor microenvironment is modified - for example after a stress-alleviation treatment - tumors characterized by different response parameters react to environmental modifications in different ways. Notably, for low cellular mechanosensitivity and low nutrient-to-mechanical response, a stress-alleviation strategy is not able to effectively reduce tumor infiltrative behavior. On the contrary, mechanosensitive tumors reduce diffuse invasion after stress-alleviation. In addition, the model highlights the influence of intrinsic tumor features, i.e. cellular diffusivity *D* and proliferation rate *r*, on the intensity of treatment response. Indeed, if on the one hand for some intrinsic proliferation rates a change in stiffness does not result in a modification of TM or IW, for the other hand for higher values of cell proliferation the two variables change significantly. These findings highlight the presence of two distinct mechanisms related to tumor progression. For a given microenvironmental stimulus, cancer cells are characterized by their sensitivity^39,40^ to the stimulus - described by the parameter *σ* in our model - and their response to it, i.e. the intensity of the phenotypic transition - described by *t*_n_ and *t*_s_. As the model suggests, there can be cases for which cell sensing is high, but cellular responses are low, and viceversa. This translates in different tumor behaviors, modulated by the intrinsic cellular diffusivity and proliferation ratio. Interestingly, these two distinct mechanisms may be targets for future therapies. For example, the model predicts that by making cells more sensitive to mechanical stimuli^41^, it is possible to elicit effective tumor responses to stress-alleviation treatments. Finally, model results highlight the existence of a trade-off between IW and TM for stress-alleviation treatments. If, on the one hand alleviation of mechanical stresses reduces tumor infiltrative tendency, on the other hand it may lead to tumors with an increased number of cells, increasing the burden in the host. Based on model results, we notice that treatments based on the alleviation of mechanical stresses in the tissue might fail for some classes of tumors, due to the complexity of the mechanisms underlying tumor progression and patient heterogeneity (e.g. different cellular mechanosensitivities, as displayed in the Results section). Patient-based estimation of intrinsic tumor features, such as cell proliferation, diffusion rates and mechanical sensitivity, would be fundamental for the design of tailored treatments for glioma patients. The experiments shown in Figure 1 represent a first step in this direction, providing some clues about the mechanical sensitivity of the cells under treatment. Indeed, even though the cell lines considered in the study are cultured in laboratory, the same assays that we described could be applied to cells deriving from specific patients. We conclude our discussion by pointing out some limitations of the current approach, as well as some possible future developments. One of the main simplifications in the model is related to the description of mechanical stresses in the tumor. In our formulation, we take into account only compressive stress states and we correlate them to cellular density^42,43^. This approach is quite phenomenological, and for a more thorough account of mechanical issues models based on continuum mechanics and mixture theory should be employed - see ref.^20,44–48^ for some recent models, and ref.^23,49^ for in-depth reviews. Nonetheless, we believe that our findings are able to provide valuable insights into the response of gliomas to microenvironmental stimuli, and to suggest new therapeutic strategies. We implicitly take into account cell apoptosis as a reduction of the net proliferation rate in the logistic growth term. Although more specific cell death mechanisms could have been considered, such as necrosis, they would impact the transient dynamics rather the long-term behavior of the system^26,38,50^. Furthermore, we modeled the migration/proliferation dichotomy in the simplest possible way, focusing on phenotypic diversity driven by nutrient levels and mechanical stresses. More complete formulations could be considered, taking into account additional tumor-related factors. As another limitation of the current approach, we carried out simulations enforcing spherical symmetry of the problem. This was to focus on the effects of model dynamics rather than on geometrical influences deriving from realistic three-dimensional geometries. As a matter of fact, these geometric features could be readily included in the model from MRI data of actual patients.

Currently, we limited the discussion to chemical and mechanical effects on tumor cells, however it is known that other intrinsic and extrinsic factors play a role in tumor progression. In fact, we plan to extend the modeling framework to investigate the interactions between gliomas and immune cells, being aware of the potential benefits of cancer immunotherapies. The dynamics of immune cells could be coupled to microenvironmental stimuli, to give a more detailed picture of glioma growth and invasion. In addition to that, the model could be used as a framework to study combined therapies in gliomas. Indeed, a decrease of invasiveness correlated to an increased mass is predicted for some tumor types as a result of stress-alleviation strategies. The model could be extended to analyze the effects of chemoterapy or radiation combined to stress-alleviation, as a mechanism to reduce tumor burden and improve the chances of cure.

We strongly believe that novel investigations on the interactions between tumor microenvironment and cellular plasticity, informed by mathematical modeling, could allow for a better understanding of disease progression, with the final goal of aiding the design of effective therapeutic treatments.

## Methods

We introduce the normalized density of proliferating and migrating glioma cells, denoted by *ρ*_p_ and *ρ*_m_, respectively. The system of equations describing the spatio-temporal evolution of the two latter quantities reads:

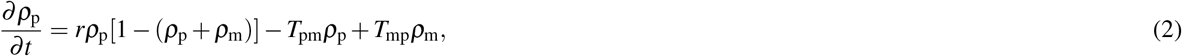

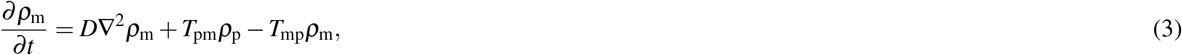

in which *T*_pm_ and *T*_mp_ are the transition rates between the two cell phenotypes. The system in (2)-(3) can be reduced to a single equation for the total density of glioma cells *ρ* = *ρ*_p_ + *ρ*_m_ by considering that *T*_pm_*ρ*_p_ = *T*_mp_*ρ*_m_. This is a plausible assumption, since the intracellular processes regulating the phenotypic switch operate at a shorter time scale than cell proliferation and migration^51^. Thus, we can express *ρ* as

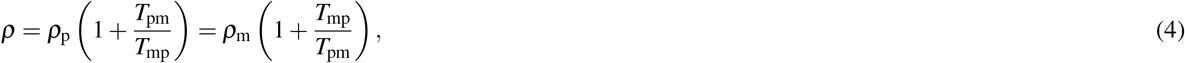

and *ρ*_p_ and *ρ*_m_ as

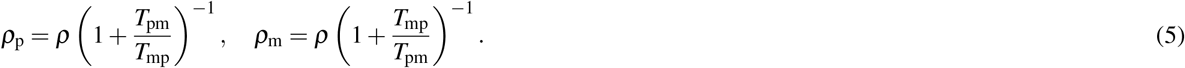

Summing equations (2) and (3), and substituting for *ρ*_p_ and *ρ*_m_ the expressions in (5), the equation for the total density *ρ* reads:

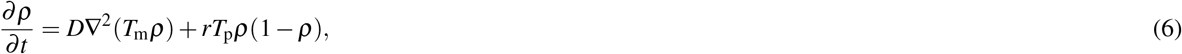

in which the quantities *T*_m_ and *T*_p_ are defined as

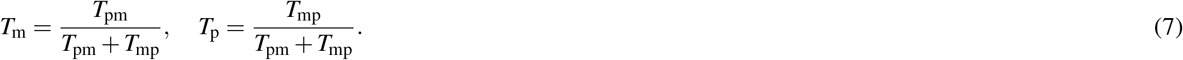

Now, we specify the equations for the normalized nutrient concentration and vascular density:

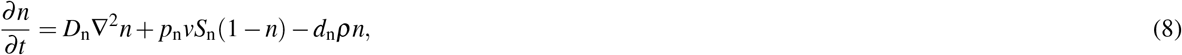

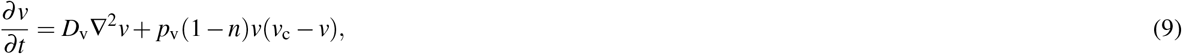

in which *D*_n_ and *D*_v_ are the diffusion coefficient of the nutrient and vascular dispersal rate, respectively; *p*_n_ is the supply rate of nutrient from the vasculature, *S*_n_ is a function accounting for the collapse of blood vessels due to mechanical compression, *d*_n_ is the nutrient consumption rate by tumor cells, *p*_v_ is the generation rate of new blood vessels and *v_c_* is the carrying capacity of blood vessels in the tumor.

To close the problem, we define suitable constitutive relations for the phenotypic transition terms, the mechanical stress sensed by tumor cells, and the vascular collapse term.

We assume that phenotypic transitions of tumor cells are driven by both chemical and mechanical effects, and we define *T*_pm_ and *T*_mp_ as

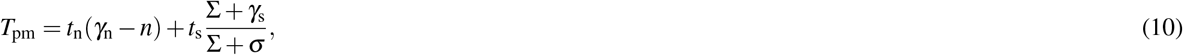

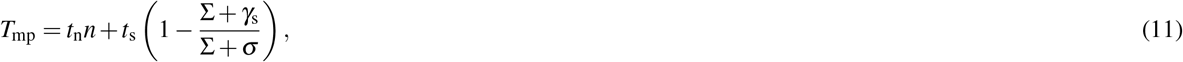

in which *t*_n_ and *t*_s_ describe the transition rates due to nutrient and mechanical stress, respectively; Σ represents the mechanical compression sensed by tumor cells, and *σ* describes cell mechanosensitivity (i.e. for high values of *σ*, cells are more refractory to transitions due to mechanical stress, and viceversa). In addition, *γ*_n_ and *γ*_s_ are small regularization terms, accounting for possible transitions when *n* or Σ are zero. For these choices of *T*_pm_ and *T*_mp_, the transition terms *T*_m_ and *T*_p_ read

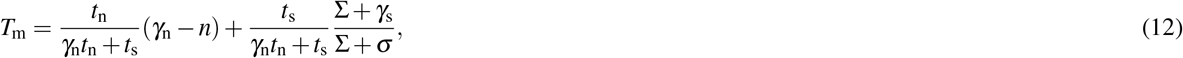

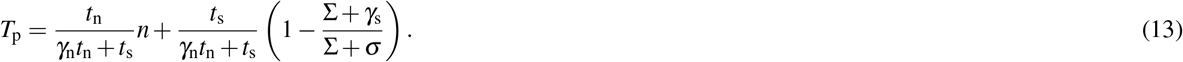

To define an expression for Σ we account for mechanical compression of the cells in a phenomenological way, as previously done for example in ref.^40,42,52^. In particular, we assume that tumor cells feel mechanical compression for increasing densities. Thus, we consider that compression is a monotonic increasing function of tumor cell density, and tends to infinity at *ρ* = 1, i.e. close to the carrying capacity. This relationship is modeled by the following expression:

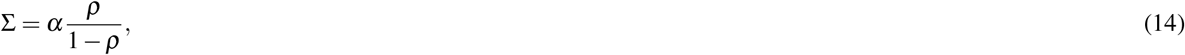

in which *α* is a constant accounting for the stiffness of the tissue.

Finally, for *S*_n_ in (8), we assume that there exists a critical mechanical compression level Σ_cr_ above which vascular functionality is compromised^36^. Since this parameter was not available in literature for gliomas, we estimate it to be of the same order of magnitude as the brain tissue stiffness. The assumed functional form accounting for this behavior is

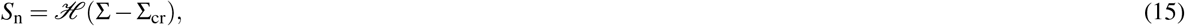

in which 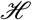 is a smooth approximation of the decreasing step function, equal to one for Σ ≈ 0Pa and equal to zero for Σ ≫ Σ_cr_.

Without loss of generality, we enforce spherical symmetry in the model equations to investigate the effects of parameter changes on a simple geometry. The spherically symmetric formulation of the problem is reported in the Appendix, whereas we summarize the parameter values used in the simulations in Table 1.

**Table 1.** Model parameters used in the simulations.

| Parameter | Description | Value | Reference |
| --- | --- | --- | --- |
| $D$ | Intrinsic diffusion coefficient of tumor cells | $2.73 \times [10^{-3}, 10^{-1}] \text{ mm}^2 \text{ d}^{-1}$ | 38 |
| $r$ | Intrinsic proliferation rate of tumor cells | $2.73 \times [10^{-4}, 10^{-2}] \text{ d}^{-1}$ | 38 |
| $\rho_c$ | Carrying density of glioma cells | $10^6 \text{ cells mm}^{-3}$ | 53 |
| $D_n$ | Diffusion coefficient of nutrient | $1.51 \times 10^2 \text{ mm}^2 \text{ d}^{-1}$ | 54 |
| $p_n$ | Nutrient supply rate from the vasculature | $10 \text{ d}^{-1}$ | 37 |
| $d_n$ | Nutrient consumption by tumor cells | $5.73 \times 10^1 \text{ d}^{-1}$ | 35 |
| $D_v$ | Vasculature dispersal rate | $5 \times 10^{-4} \text{ mm}^2 \text{ d}^{-1}$ | 55 |
| $p_v$ | Vasculature formation rate | $10^{-1} \text{ d}^{-1}$ | 56 |
| $v_c$ | Vasculature carrying capacity | 3 | 36 |
| $\Sigma_{cr}$ | Critical stress for vasculature collapse | $10^2 \text{ Pa}$ | estimated |
| $\gamma_n$ | Regularization term in (10) | 1.01 | model specific |
| $\gamma_s$ | Regularization term in (11) | $10^{-2} \sigma \text{ Pa}$ | model specific |

### Experimental procedure

Cell culture. Human glioma cell lines, H4 and A172, were obtained from American Type Culture Collection (ATCC) and were maintained in Dulbecco ‘s Modified Eagle ‘s Medium (DMEM) supplemented with 10% Fetal Bovine Serum (FBS) and 1% antibiotics. Both cell lines were incubated at 37°C and 5% CO_2_ in a humidified incubator.

#### Application of mechanical compression

For the generation of solid stress in cancer cells, the transmembrane pressure device was employed as previously described^57^. Briefly, cancer cells were seeded in the inner chamber of a transwell insert (Greiner Bio-one), and a 1.5mm-thick agarose gel was added on top of the cells preventing any contact between piston and cells, providing a uniform distribution of the applied force. A piston of a predefined weight was placed on the top of the agarose gel and the cells were exposed to 4mmHg stress. Control cells were covered with an agarose gel.

Proliferation Assay. Cancer cells were seeded in the inner chamber of a transwell insert (10^5^ cells/insert) overnight. Alamar blue reagent (Thermo) was subsequently added in culture medium (10% v/v) and cell number was calculated prior- and post-compression according to the manufacturer ‘s instructions. Control cells were covered with an agarose cushion only.

Wound Closure assay. A wound healing assay was performed on cancer cells as previously described^57^. In brief, cancer cells were seeded in the inner chamber of a transwell insert (2 × 10^4^ cells/insert) and allowed to form a monolayer. A scratch wound was then introduced on the cell monolayer, cells were washed twice with PBS and then were stimulated with 4mmHg stress for 16h. Control cells were covered with an agarose gel (i.e. 0mmHg). Images from at least three different fields from each condition were taken at 0h and 16h. The cell-free area was quantified using the ImageJ software^58^. Quantification was performed for each condition using the following formula: (Area of the wound at 0h – Area of the wound at 16h) / (Area of the wound at 0h).

## Acknowledgements

MMH is supported by Measures for the Establishment of Systems Medicine projects SYSIMIT (01ZX1308D) and Sys-Stomach (01ZX1310C) by the Federal Ministry of Education and Research (BMBF), Germany, and by iMed - the Helmholtz Initiative on Personalized Medicine. HH and PM are supported by Mic2Mode-I2T (01ZX1710B) and HH is supported by SYSIMIT (01ZX1308D) and MulticellML (01ZX1707C) by the Federal Ministry of Education and Research (BMBF) and by the SYSMIFTA (031L0085B) of the ERACOSYSMED initiative. MK and TS acknowledge the funding support of the European Research Council Starting Grant (ERC-2013-StG-336839 ReEngineeringCancer).

## Competing interests

The authors declare no competing financial interests.

## Appendix

The final system of equations in spherical symmetry reads:

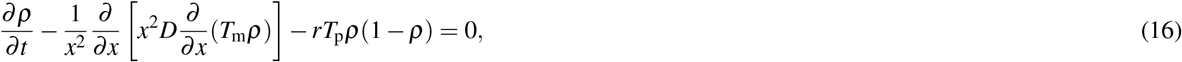

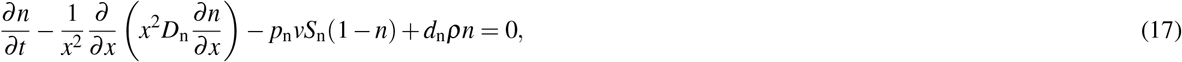

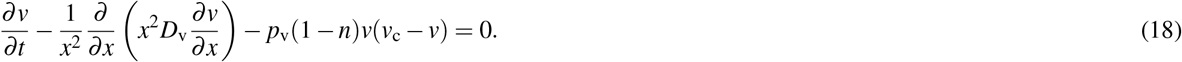

In which *x* is the radial coordinate. To close the problem, we define the following boundary and initial conditions:

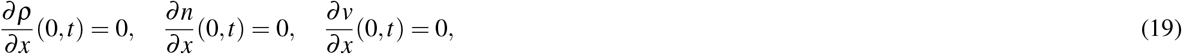

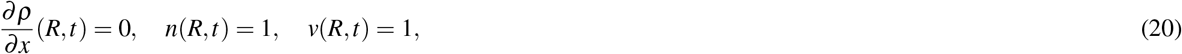

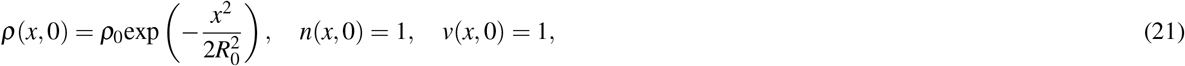

in which *R* is the external radius of the domain, *ρ*_0_ is the initial tumor density and *R*_0_ is the initial radius of the tumor. A schematic of the model is shown in Figure A1.

Regarding the model parameters, these are taken from published data whenever available, or estimated by suitable physical and biological arguments. Table 1 summarizes the parameter values, together with their units and reference sources. We simulate the growth of a tumor over 1 year, and consider the following parameter values: *R* = 100mm, *R*_0_ = 2mm, *ρ*_0_ = 0.1.

Once the mathematical model is parametrized, we proceed to its spatio-temporal discretization. This is performed via the finite element method, using linear elements; for time discretization, we use a standard backward Euler scheme, an implicit method. Once the equations are discretized, the model is implemented using the open source software FEniCS^59^.

**Figure A1.**
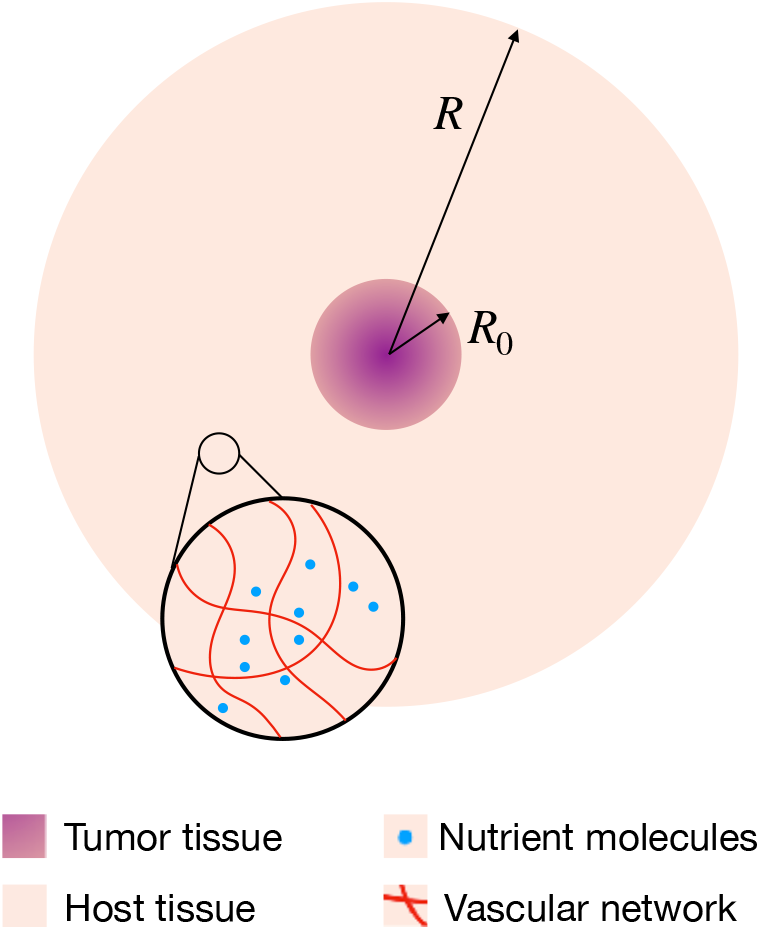
Schematics of the mathematical model. The tumor is placed at the center of the host tissue, and the inlet highlights the presence of the vascular network and nutrient molecules in the system. *R*_0_ and *R* denote the initial tumor radius and the computational domain, respectively.

## Supplementary Materials

**Figure S1.**
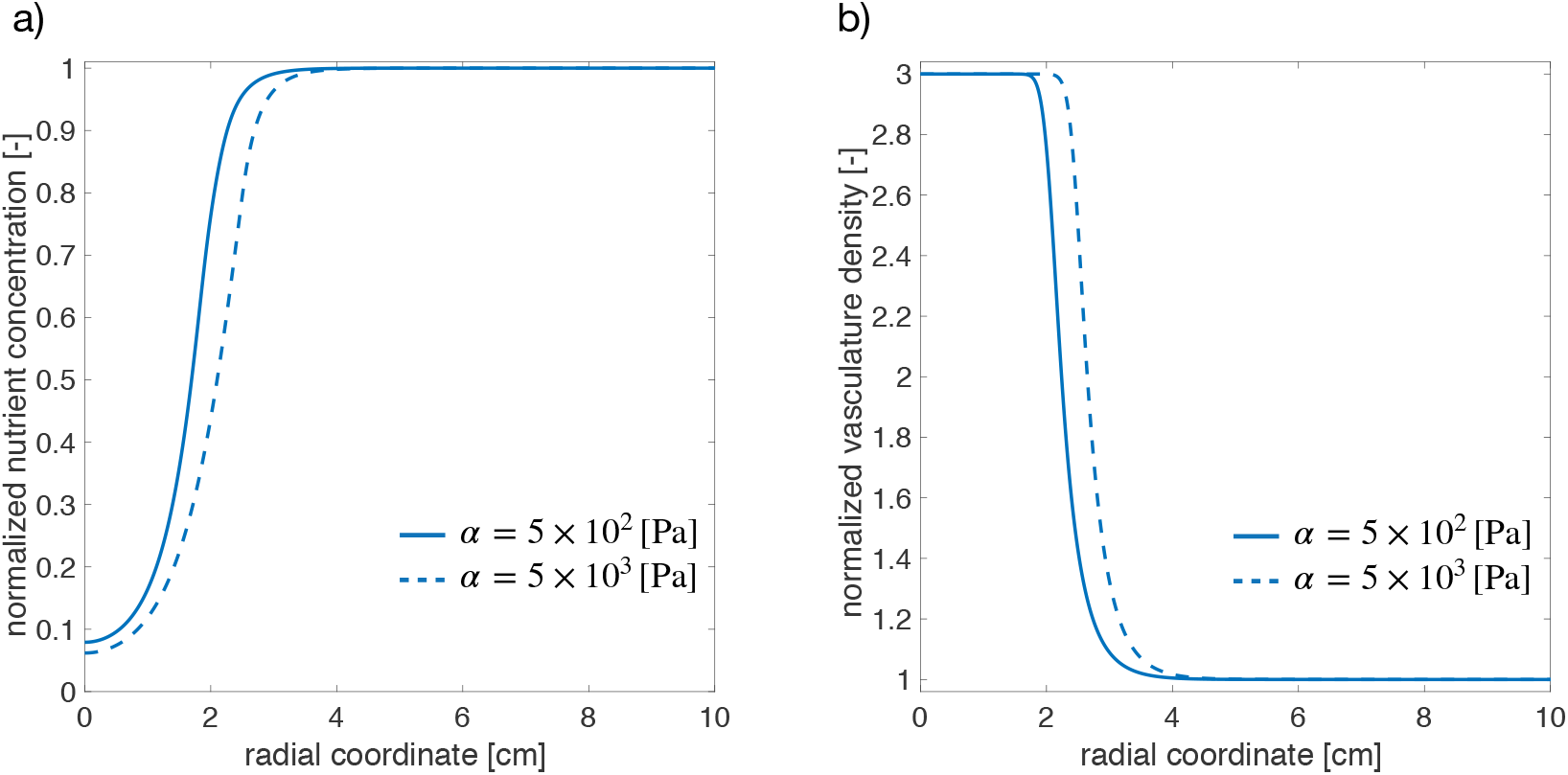
Change in nutrient concentration (a) and vasculature density (b) after a stress-alleviation treatment (*α* is reduced from 5 × 10^3^Pa to 5 × 10^2^Pa). The simulations refer to the case of *D* = 2.73 × 10^−1^mm^2^d^−1^ and *r* = 2.73 × 10^−2^d^−1^.

**Figure S2.**
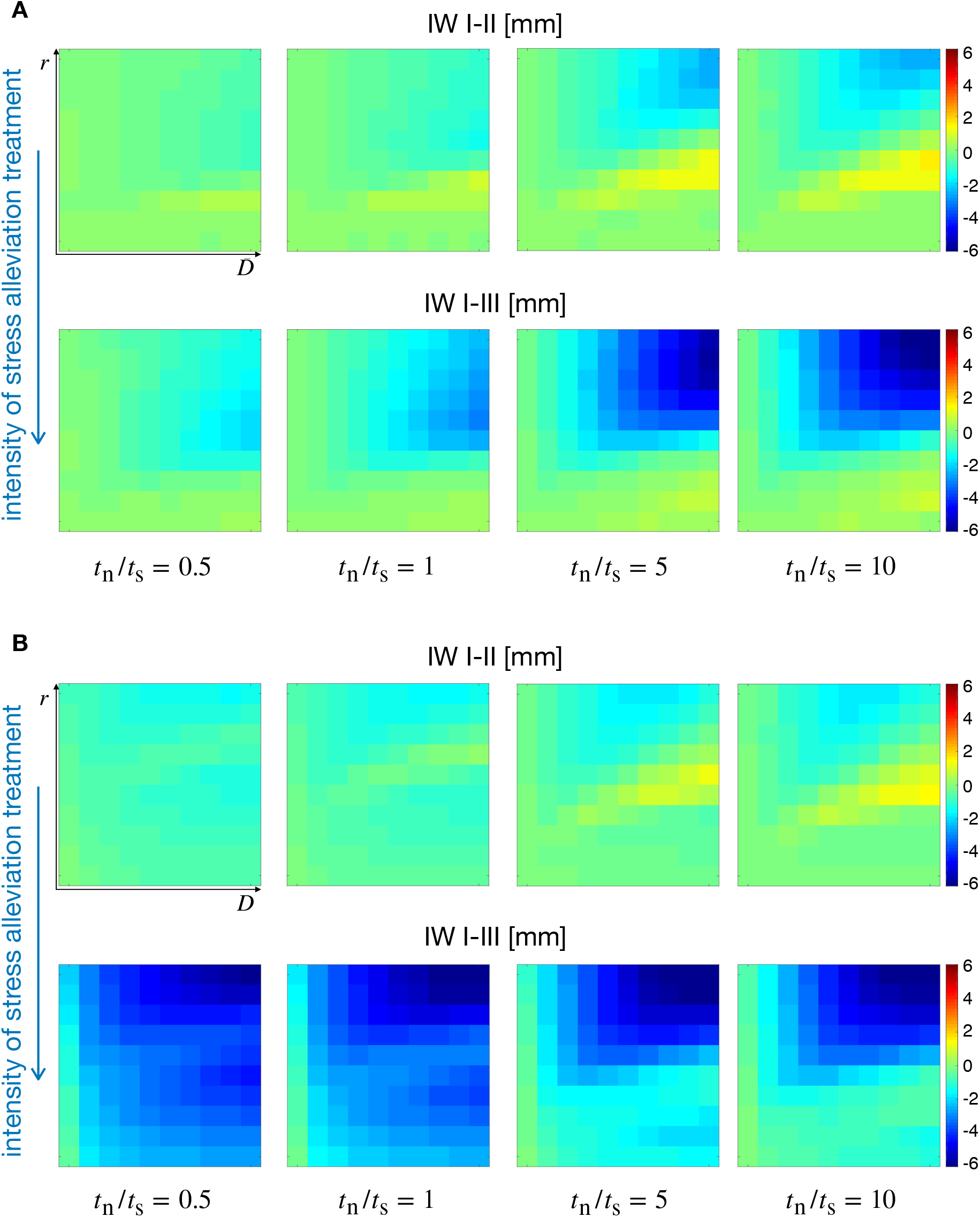
Simulation maps displaying the impact of chemo-mechanically induced transitions on tumor IW. In both cases (**A, B**), the top row shows the IW difference when tissue stiffness varies from *α* = 10^3^Pa to *α* = 5 × 10^2^Pa, whereas the bottom row displays the IW variations for *α* = 5 × 10^3^Pa to *α* = 5 × 10^2^Pa. Simulations were obtained for low, i.e. *ασ*^−1^ = [10^−2^,10^−1^] (**A**), and high, i.e. *ασ*^−1^ = [10^1^,10^2^] (**B**) mechanosensitivity.

**Figure S3.**
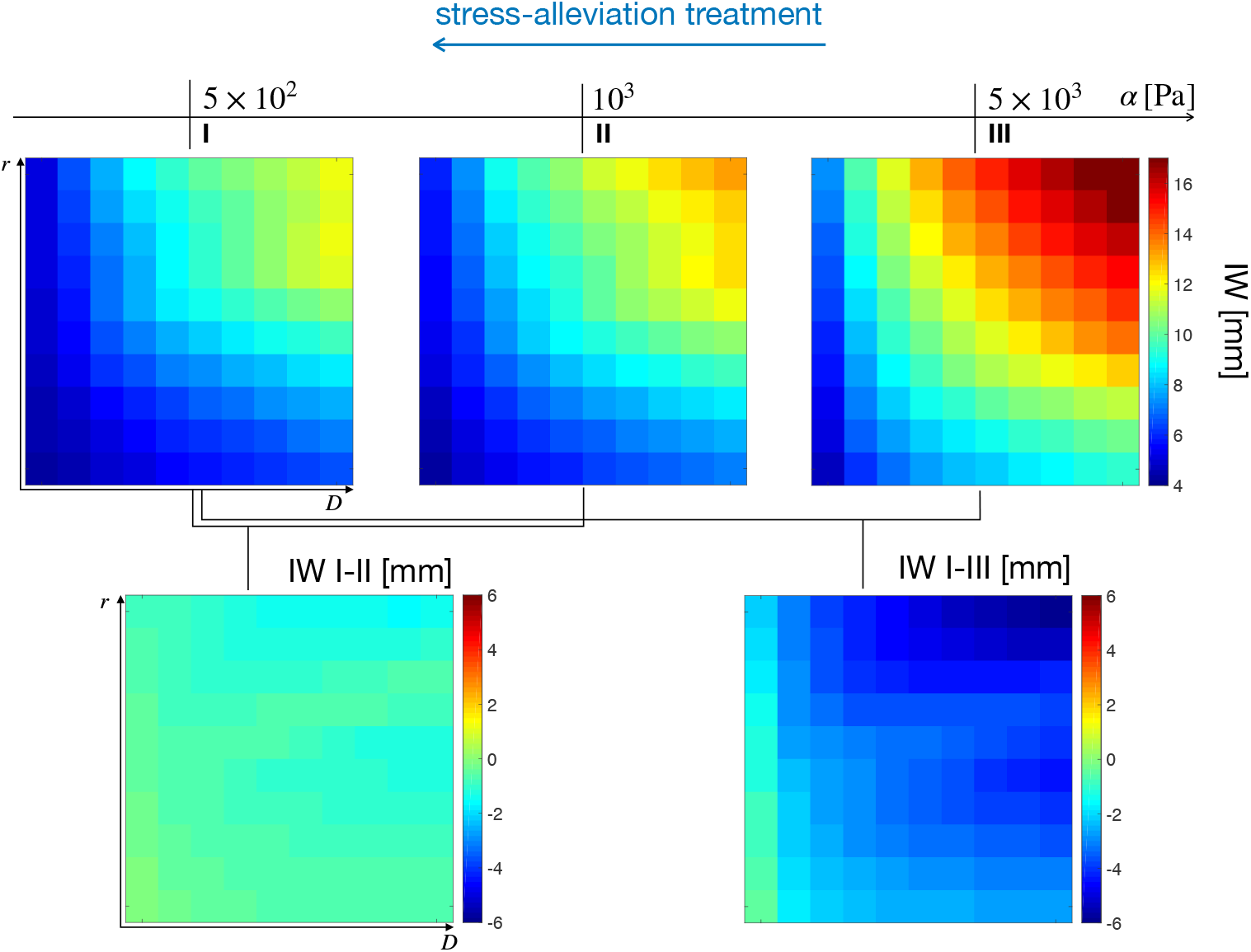
Simulation maps displaying the effects of chemo-mechanically induced transitions on tumor IW. The top row shows three IW maps for different values of *α*, whereas the bottom row displays the IW variation occurring at the different stiffness points. For these simulations, we used *t*_n_/*t*_s_ = 0.5 and *ασ*^−1^ = [10^1^,10^2^].

**Figure S4.**
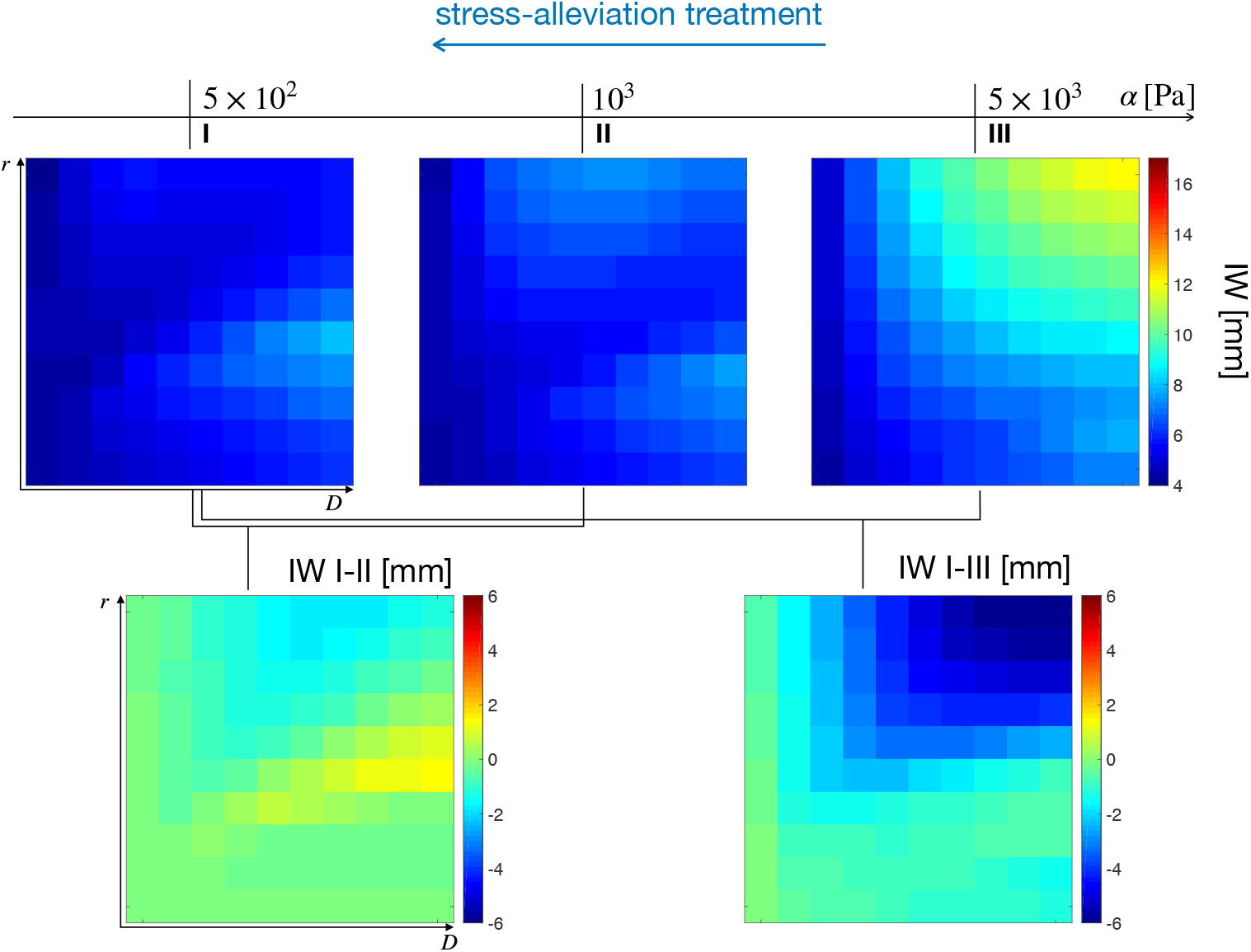
Simulation maps displaying the effects of chemo-mechanically induced transitions on tumor IW. The top row shows three IW maps for different values of *α*, whereas the bottom row displays the IW variation occurring at the different stiffness points. For these simulations, we used *t*_n_/*t*_s_ = 10 and *ασ*^−1^ = [10^1^,10^2^].

**Figure S5.**
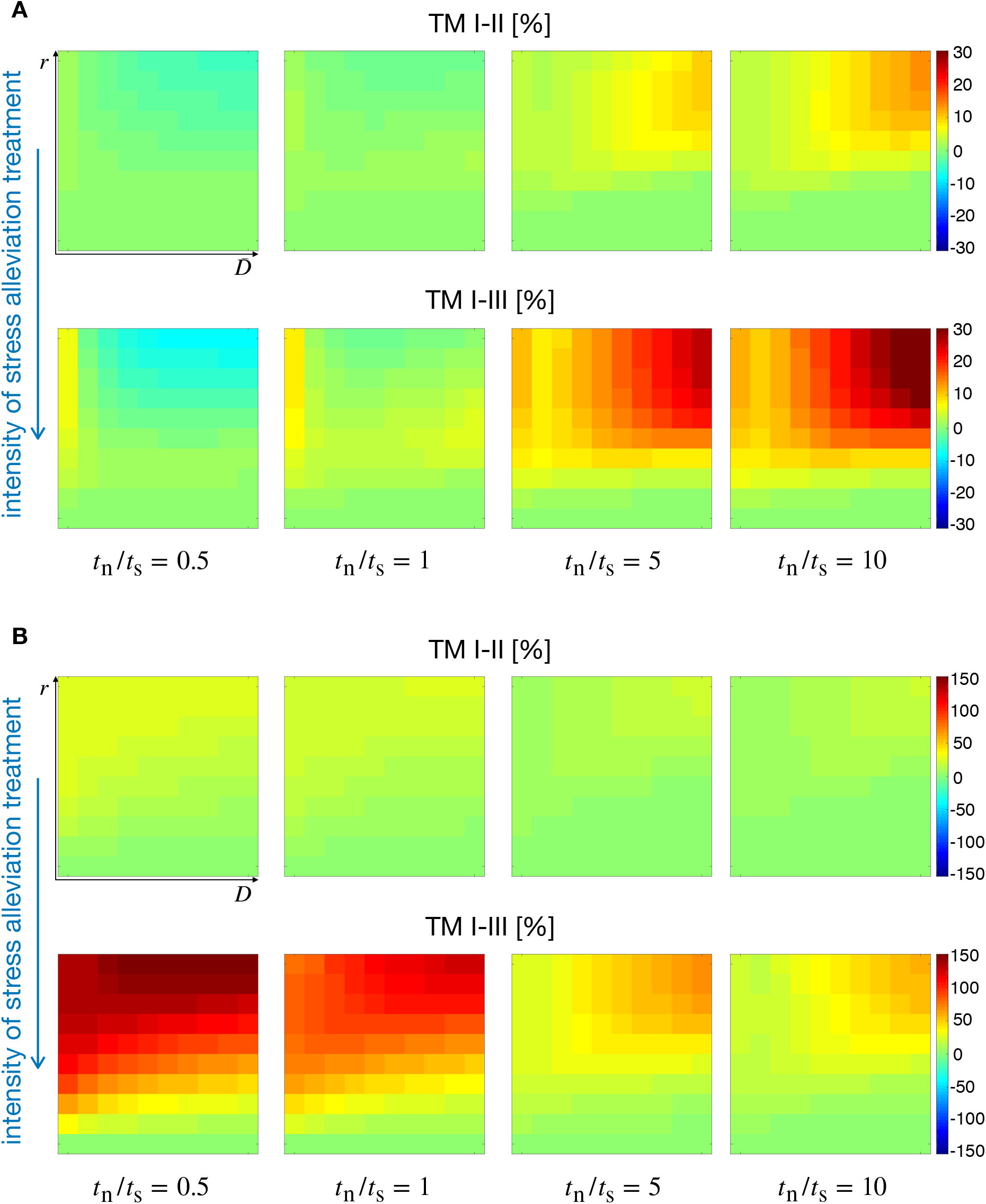
Simulation maps displaying the impact of chemo-mechanically induced transitions on TM. In both cases (**A, B**), the top row shows the TM difference for a reduction in tissue stiffness from *α* = 10^3^Pa to *α* = 5 × 10^2^Pa, whereas the bottom row displays the TM variations for *α* = 5 × 10^3^Pa to *α* = 5 × 10^2^Pa. Simulations were obtained for low, i.e. *ασ*^−1^ = [10^−2^,10^−1^] (**A**), and high, i.e. *ασ*^−1^ = [10^1^,10^2^] (**B**) mechanosensitivity.

**Figure S6.**
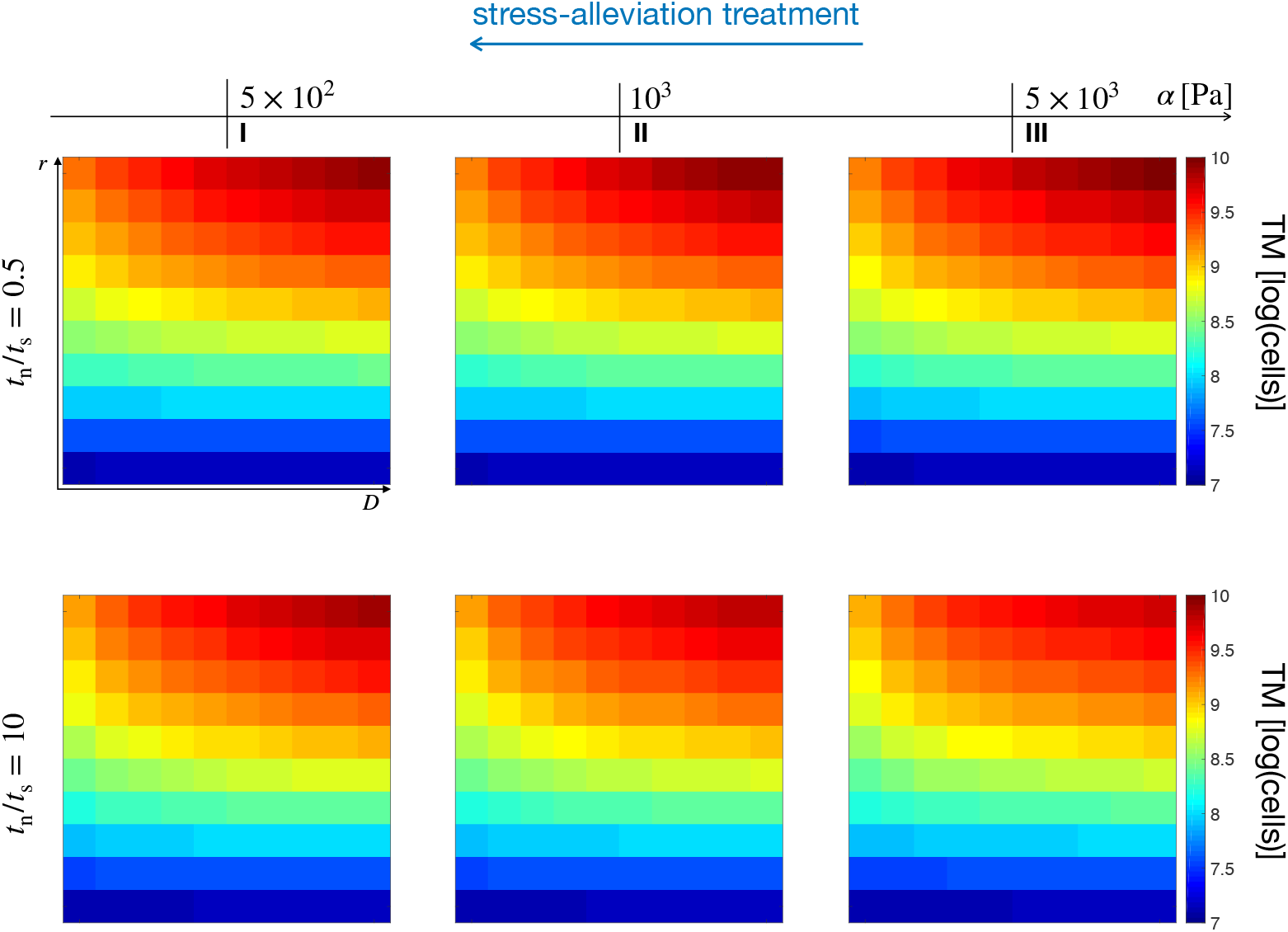
Simulation maps displaying the effects of chemo-mechanically induced transitions on TM. The top row shows three TM maps for different values of *α* at the ratio *t*_n_/*t*_s_ = 0.5, whereas the bottom row displays TM values over the (*D, r*) space for the different stiffness points at the *t*_n_/*t*_s_ = 10 ratio. The simulations refer to the low mechanosensitivity case, i.e. *ασ*^−1^ = [10^−2^,10^−1^].

**Figure S7.**
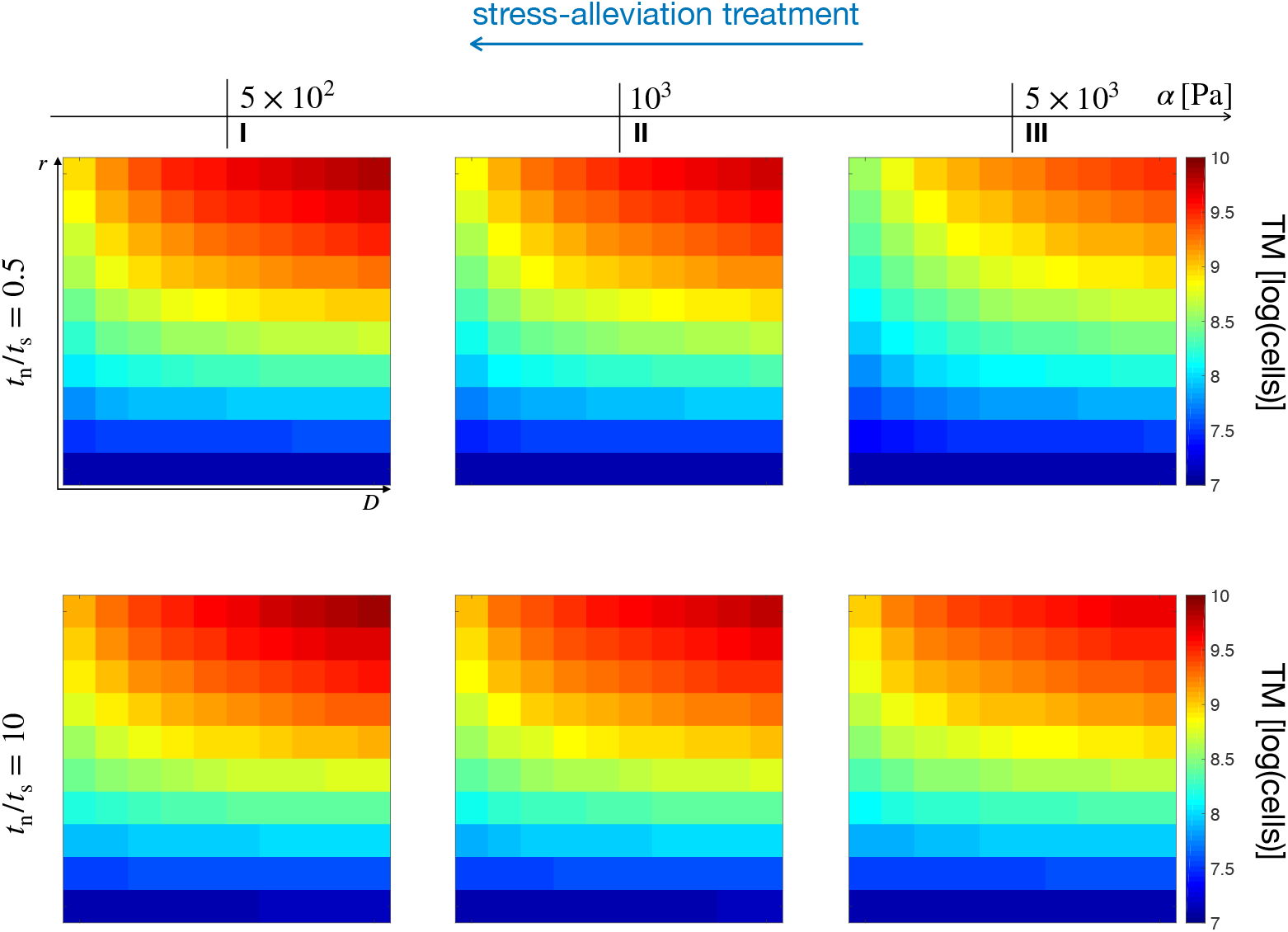
Simulation maps displaying the effects of chemo-mechanically induced transitions on TM. The top row shows three TM maps for different values of *α* at the ratio *t*_n_/*t*_s_ = 0.5, whereas the bottom row displays TM values over the (*D, r*) space for the different stiffness points at the *t*_n_/*t*_s_ = 10 ratio. The simulations refer to the high mechanosensitivity case, i.e. *ασ*^−1^ = [10^1^,10^2^].

